# Adaptive Laboratory Evolution under gradual ciprofloxacin stress leads to efflux-mediated ciprofloxacin resistance and an increase in endogenous antibiotic production in *Streptomyces coelicolor*

**DOI:** 10.64898/2025.12.22.695978

**Authors:** Ankita Nag, Charushila Kumbhar, Sarika Mehra

## Abstract

*Streptomyces* are soil-dwelling bacteria harboring a vast reservoir of transcriptionally silent antibiotic-resistant genes. Mobilization of these resistance genes to pathogens results in the dissemination of multidrug resistance when exposed to differential antibiotic pressure. To determine the evolution of resistance, we employed adaptive laboratory evolution (ALE) in *Streptomyces coelicolor* under gradually increasing concentrations of ciprofloxacin, starting at the minimum inhibitory concentration (MIC) to 4X MIC. The evolved strain CipRr exhibited elevated resistance towards ciprofloxacin with cross-resistance to other fluoroquinolones, such as ofloxacin and moxifloxacin, and collateral sensitivity to the protein synthesis inhibitors chloramphenicol and tetracycline. Compared to the WT cells, CipRr showed elevated production of secondary metabolites actinorhodin and undecylprodigiosin, altered morphology, and reduced ROS production. Whole Genome Sequencing revealed mutations in transcriptional regulators and other metabolic genes, leading to enhanced activity of efflux pumps that conferred high-level ciprofloxacin resistance. Overall, this study demonstrates that evolutionary strategies in antibiotic producers such as streptomyces can help uncover the genetic determinants linked to antibiotic resistance, secondary metabolism, and morphological differentiation.

**Importance:** This study employs adaptive laboratory evolution in the antibiotic-producing bacterium Streptomyces coelicolor to investigate how exposure to gradually increasing concentrations of the fluoroquinolone ciprofloxacin drives the emergence of high-level resistance and cross-resistance to related drugs, while simultaneously resulting in collateral sensitivity towards other antimicrobial agents. This study also highlights that the evolution of high-level resistance towards ciprofloxacin significantly alters the production of secondary metabolites and cellular morphology. Furthermore, we identify mutations in regulatory and metabolic genes that result in enhanced efflux pump activation, advancing our understanding of the molecular mechanisms underlying high-level resistance. This study thus highlights the complex interplay between antibiotic resistance, metabolism, and morphological development.

## Introduction

*Streptomyces* are soil-dwelling bacteria that produce 60% of the world’s total antibiotics. These antibiotic producers in turn, have evolved a self-resistance mechanism that protects these bacteria from their own antibiotics (1, 2). Furthermore, various *Streptomyces* collected from different geographical areas show elevated resistance to a wide group of drugs, implicating their role as a potential reservoir of antibiotic resistance genes (ARGs) (3). Mobilization of these resistance genes to clinical pathogens is a crucial factor in the dissemination of antimicrobial resistance (AMR) in the environment (4). The synthesis of antibiotics by *Streptomyces* is dependent on a set of genes termed the Biosynthetic Gene Cluster (BGCs) (5). Interestingly, the resistance genes for these endogenous antibiotics are co-located in the same gene cluster. Bioinformatics studies reveal that *Streptomyces* harbor a wide array of BGCs, which remain cryptic in nature with low or no expression at all. Many of these resistant genes find their way through Horizontal Gene Transfer (HGT) into clinical pathogens, thus spreading AMR (6).

Given the presence of many cryptic resistance gene expressions, innovative techniques such as adaptive laboratory evolution (ALE) are required to identify latent resistance mechanisms (7). ALE is predominantly used to understand the adaptive potential of microorganisms in biotechnology (8). It has been adopted to derive the mechanisms behind drug resistance in various microbial pathogens (9). This method relies on exposing the bacteria to gradually increasing antimicrobial stress at different time periods, revealing the role of potential cryptic genes involved in resistance (10). Evolutionary studies on *Streptomyces coelicolor* have reported that AMR in nature can arise and persist even in the presence of sub-lethal concentrations of antibiotics, as antibiotics continue to work as selective weapons rather than just harmless signals (11). Additionally, previous transcriptomics studies have documented a range of resistance determinants that may emerge in *S. coelicolor* when subjected to shorter exposure times to ciprofloxacin (12, 13).

In the present study, we utilize ALE to determine the resistance mechanism in *S. coelicolor.* We subjected the bacteria to increasing concentrations of ciprofloxacin, a fluoroquinolone, starting at the Minimum Inhibitory Concentration (MIC) with a gradual increase to 4X MIC and assessed the accumulated mutations that modulate antibiotic resistance, secondary metabolism, and morphological development.

## Results

### CipRr displays altered susceptibility towards different classes of drugs

When compared to WT, CipRr displayed a 15-fold increase in resistance towards ciprofloxacin. Cross-resistance was also observed to other fluoroquinolones with 12-fold and 6-fold increase in MIC against ofloxacin and moxifloxacin, respectively (Table 1). In contrast, CipRr exhibited increased susceptibility to the ribosome-binding drugs chloramphenicol and tetracycline and the DNA gyrase inhibitor novobiocin, with a 2-fold reduction in MIC relative to WT. Previous studies have shown that resistance to one fluoroquinolone results in the development of cross-resistance to other fluoroquinolones (14). Besides, resistance towards fluoroquinolones also results in collateral sensitivity towards other antibiotics (15). On similar lines, our findings indicate that prolonged ciprofloxacin exposure in *S. coelicolor* not only provides enhanced resistance to fluoroquinolones but also increases susceptibility towards other non-fluoroquinolone drugs.

**Table 1:** Determination of Minimum Inhibitory Concentration (MIC) of WT and CipRr towards multiple antibiotics.

| Antibiotics | Minimum Inhibition Concentration (MIC) |  |
| --- | --- | --- |
|  | (µg/ml) |  |
|  | WT | CipRr |
| Ciprofloxacin | 80 | 1200 |
| Ofloxacin | 40 | 480 |
| Moxifloxacin | 80 | 480 |
| Chloramphenicol | 50 | 25 |
| Tetracycline | 50 | 25 |
| Ampicillin | 50 | 50 |
| Novobiocin | 5 | 2.5 |
| Spectinomycin | 12.5 | 12.5 |
| Oleandomycin | 25 | 25 |

### CipRr displays distinct phenotypic morphology as compared to WT *S. coelicolor* cells

Our results indicate that the CipRr mutant exhibits differential susceptibility to a range of structurally diverse drugs (Table 1). To further investigate whether the resistance to fluoroquinolones in CipRr is a result of any alteration phenotype, we analyzed the morphology of CipRr mutants using Cryo-SEM imaging and compared them to the WT. A distinct morphology with nodular-like mycelia was observed in CipRr, which was not seen in WT cells (Figure 1a). Such morphological changes have been previously linked to alterations in cell wall composition that facilitate survival under antibiotic stress (16, 17). We therefore hypothesize that CipRr may have adopted an altered phenotype to survive increasing antibiotic pressure and provide high-level resistance towards them.

**Figure 1:**
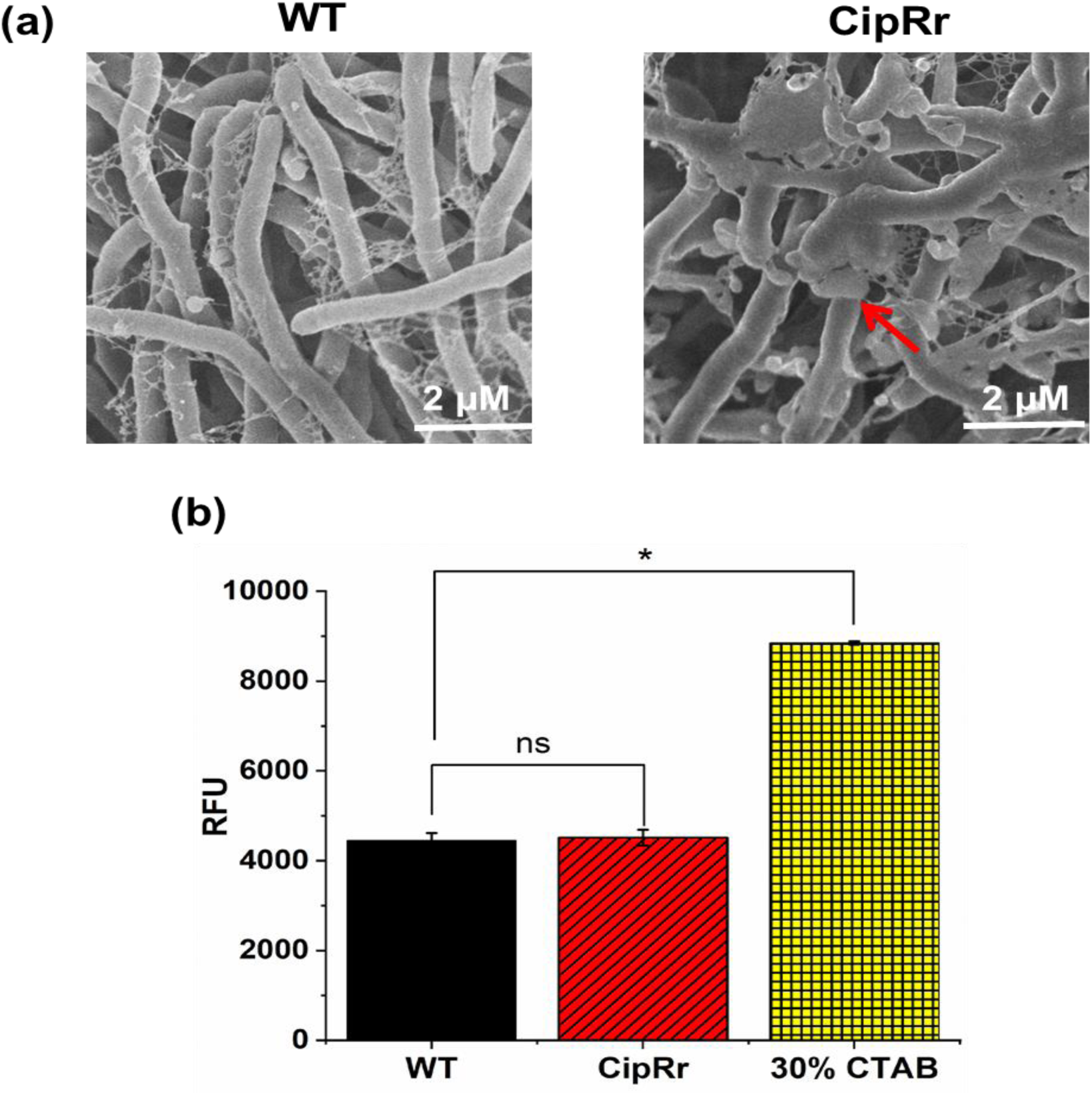
**Comparison of cell morphology and membrane integrity in *S. coelicolor* WT and CipRr strains**. (a) Cryo-SEM imaging of *S. coelicolor* strain M145 and ciprofloxacin-resistant CipRr strains. Cells were grown till 1 O.D, washed with 1X PBS followed by 10% ethanol to remove the media contaminants before imaging under the Cryo-FEG SEM (JSM-7600F) platform. Scale bars correspond to 2 µM. Red arrow indicates the nodular-like morphology seen in CipRr strains. (b) Cell membrane integrity measured using NPN assay. WT cells treated with 30% CTAB were taken as the positive control. The assay was performed in triplicates and statistical significance was calculated with respect to WT cells. * indicates P-value 0.05; ns indicates no significance.

To determine if the altered membrane permeability also contributes to the differential resistance of CipRr towards different drugs, we monitored their permeability using the NPN assay as described in the methods. NPN cannot penetrate an intact membrane; however, a change in membrane permeability leads to increased binding of the dye to the hydrophobic membrane, thereby leading to increased fluorescence(18). We found that despite a distinct morphology, the NPN binding profile in the CipRr strain was the same as that of WT cells (Figure 1b), indicating no significant change in permeability. The cells treated with 30% CTAB showed increased fluorescence and served as a positive control for disrupted membrane integrity.

### Increased production of actinorhodin and undecylprodigiosin seen in CipRr strains

A previous study in our lab has shown that brief exposure to ciprofloxacin results in differential production of endogenous antibiotics in *S. coelicolor* under different growth conditions (12, 13). To assess the impact of ciprofloxacin adaptation on secondary metabolite production in *S. coelicolor*, we quantified two endogenous antibiotics, actinorhodin and undecylprodigiosin yields in CipRr strains and compared to the WT. Both antibiotics exhibited significantly higher time-dependent accumulation in CipRr as compared to WT (Figure 2a). At the end of 96 hours, actinorhodin levels increased to 5-fold, whereas undecylprodigiosin increased by nearly 2-fold in CipRr (Figure 2b and 2c).

**Figure 2:**
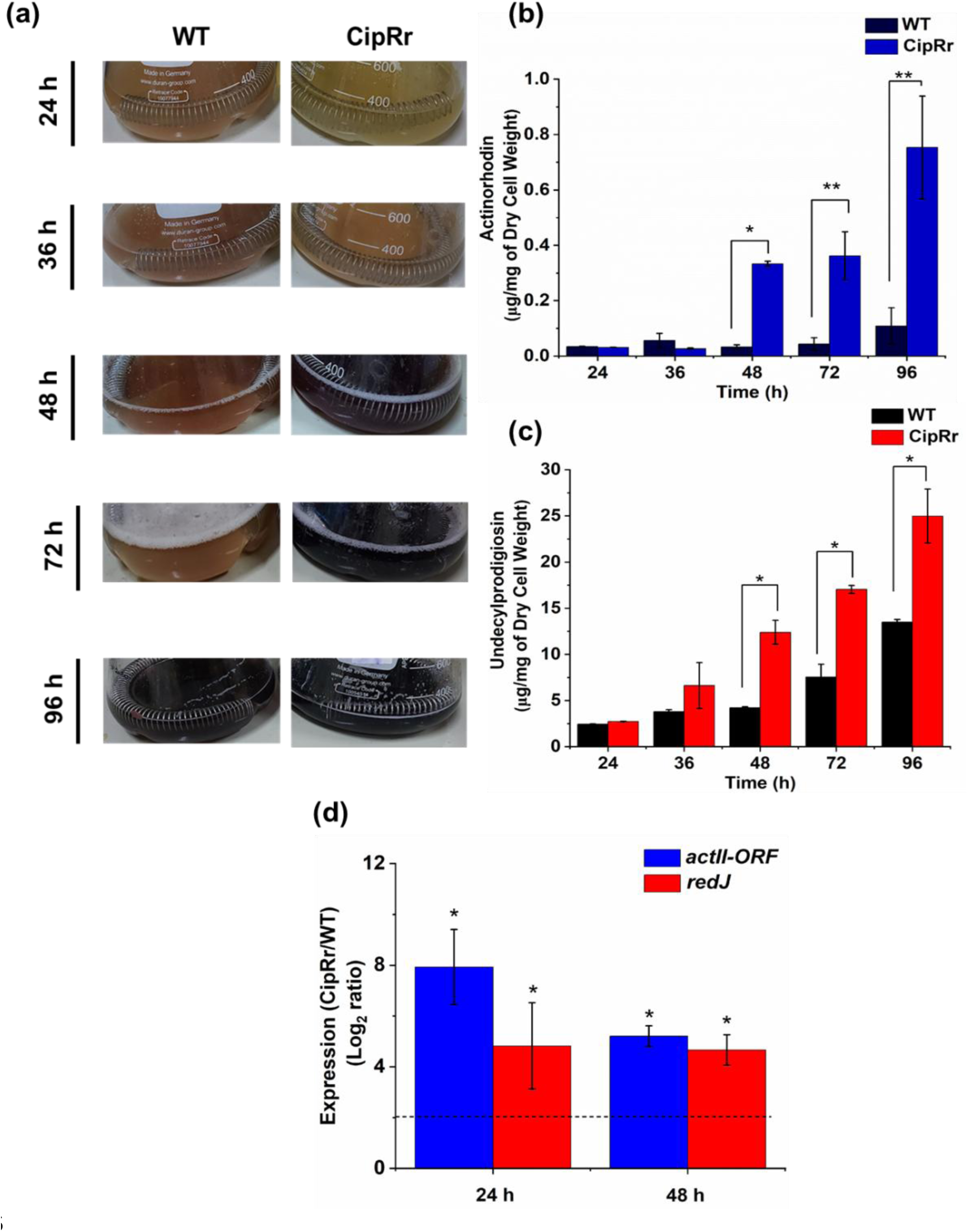
**Estimation of antibiotic production in *S. coelicolor* WT and CipRr cells**. (a) Antibiotic production observed in WT and CipRr cultures grown in R5 media. Samples were taken at specific time intervals of 24, 36, 48, 72 and 96 h respectively and imaged. Blue corresponds to actinorhodin, and red corresponds to undecylprodigiosin. (b) Quantitative estimation of actinorhodin in WT and CipRr cells determined at 24, 36, 48, 72 and 96 hours respectively. (c) Quantitative estimation of undecylprodigiosin in WT and CipRr measured at 24, 36, 48, 72 and 96 h respectively. (d) Expression of *actII-orf* and *redj* measured in WT and CipRr strains using Real Time Quantitative PCR. Samples were taken at 24 h and 48 h respectively and expression was normalized with 23S rRNA as the house keeping gene followed by normalization with the WT cells taken at the specified intervals. Statistical significance calculated with respect to WT cells. ** indicates p-value <0.005, * indicates p-value <0.05, ns indicates no significance.

We further analyzed the expression profile of *actII-ORF,* which is involved in the export of actinorhodin and *redJ* genes, required for the production of undecylprodigiosin, using qRT-PCR. In CipRr, both *actII-orf* and *red*J genes were upregulated to 8-fold (log_2_ ratio) and 5-fold (log_2_ ratio), respectively, at 24 h compared to WT cells (Figure 2d). Similarly, at 48 h, both *actII-ORF* levels and *redJ* remained upregulated to nearly 4-fold (log_2_ ratio) (Figure 2d). We found this data to be consistent with the antibiotic production profile seen in CipRr strains (Figure 2a). Previous studies have reported that physiological and chemical stress stimulate secondary metabolites in *Streptomyces* (19, 20). Our results suggest that prolonged exposure to ciprofloxacin results in fixation of mutations that enable CipRr to produce significantly higher yields of the endogenous antibiotics.

### CipRr displays increased tolerance to oxidative stress

Bacterial exposure to bacteriostatic drugs such as ciprofloxacin is known to induce oxidative stress by increasing the intracellular reactive oxygen species (ROS). In response, the bacteria accumulate mutations that facilitate survival (21, 22). We observed elevated production of actinorhodin in CipRr as compared to the WT strain (Figure 3a). Previous studies have identified actinorhodin as a redox-active antibiotic that induces the expression of genes responsible for quenching ROS stress (23). We further screened the susceptibility of CipRr towards different oxidants. Similar susceptibility towards the oxidant H_2_O_2_ was observed for both CipRr and WT (Figure 3a). In contrast, CipRr displayed increased resistance to the strong oxidant HOCl (Figure 3b). Quantification of ROS using the DCFDA assay demonstrated that CipRr generated nearly 5-fold lower intracellular ROS than WT (Figure 3c). This data suggests that adaptive evolution under ciprofloxacin exposure confers enhanced oxidative stress tolerance.

**Figure 3:**
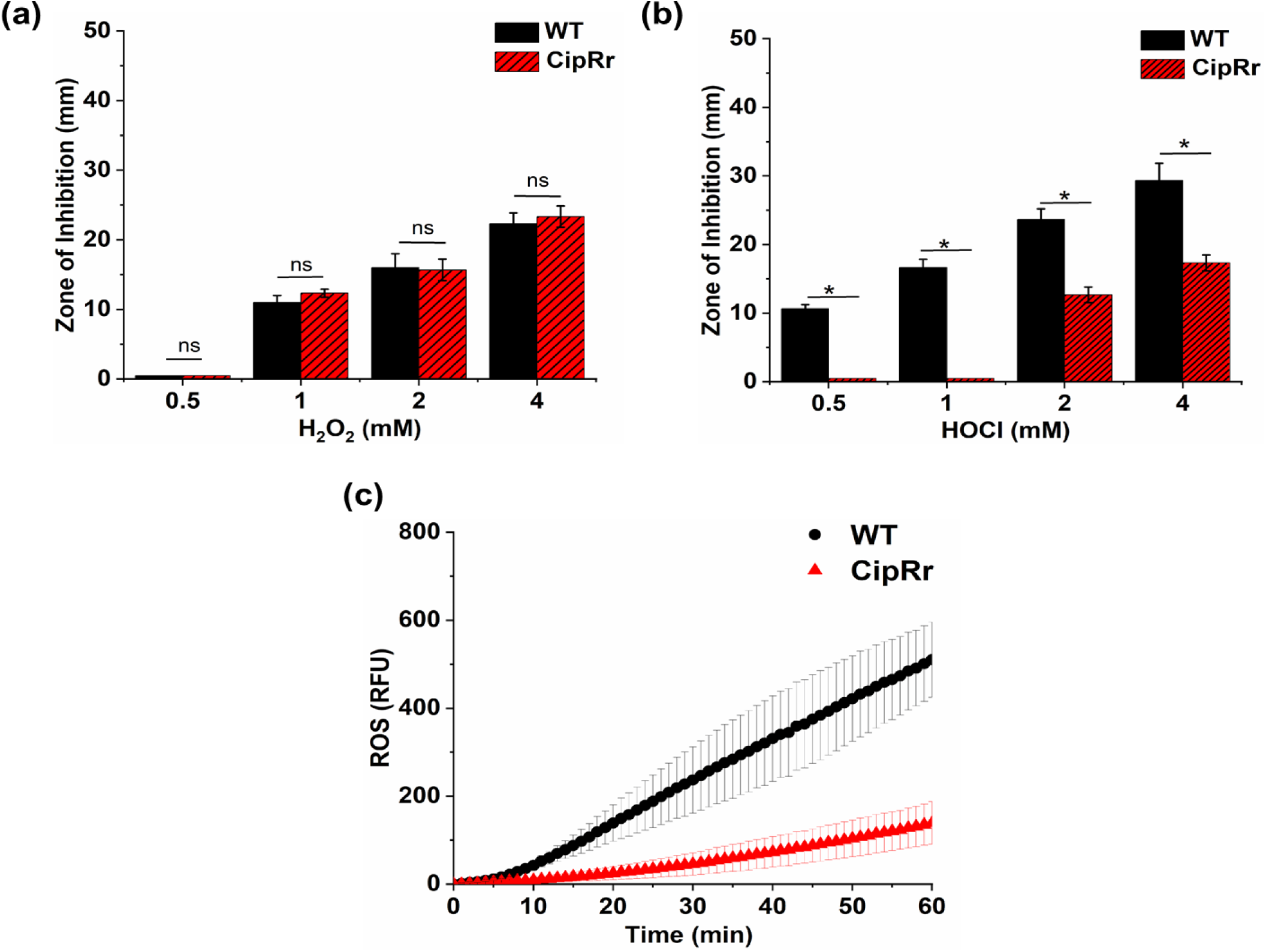
**Determination of oxidative stress tolerance in WT and CipRr strains**. Susceptibility of WT and CipRr measured towards (a) H_2_O_2_ and (b) HOCl. Disc Diffusion assay was used to measure susceptibility towards oxidants. Zone sizes are measured in millimeters. Statistical significance is calculated with respect to WT cells. (c) Measurement of intracellular levels of ROS in WT and CipRr was measured using the DCFDA assay. DCFDA at a concentration of 10 µM was used for the assay. Error bars were calculated based on three biological replicates. * Indicates p<0.05, ns indicates no significance.

### Overexpression of efflux pumps is a probable factor of resistance in CipRr strains

To understand the role of efflux pumps in ciprofloxacin resistance, susceptibility assays were conducted with CipRr strains in the presence of efflux pump inhibitors. We found that neither Carbonyl cyanide m-chlorophenyl hydrazine (CCCP), which targets MFS transporters, nor Sodium Orthovanadate (NaOVa), which targets ABC transporters alone, restored the ciprofloxacin susceptibility in CipRr. However, a combination of both inhibitors at sub-inhibitory concentrations partially restored the MIC in CipRr strains (Figure 4a). This data indicates that elevated resistance in CipRr is partially contributed to by overexpression of efflux pumps belonging to multiple families.

**Figure 4:**
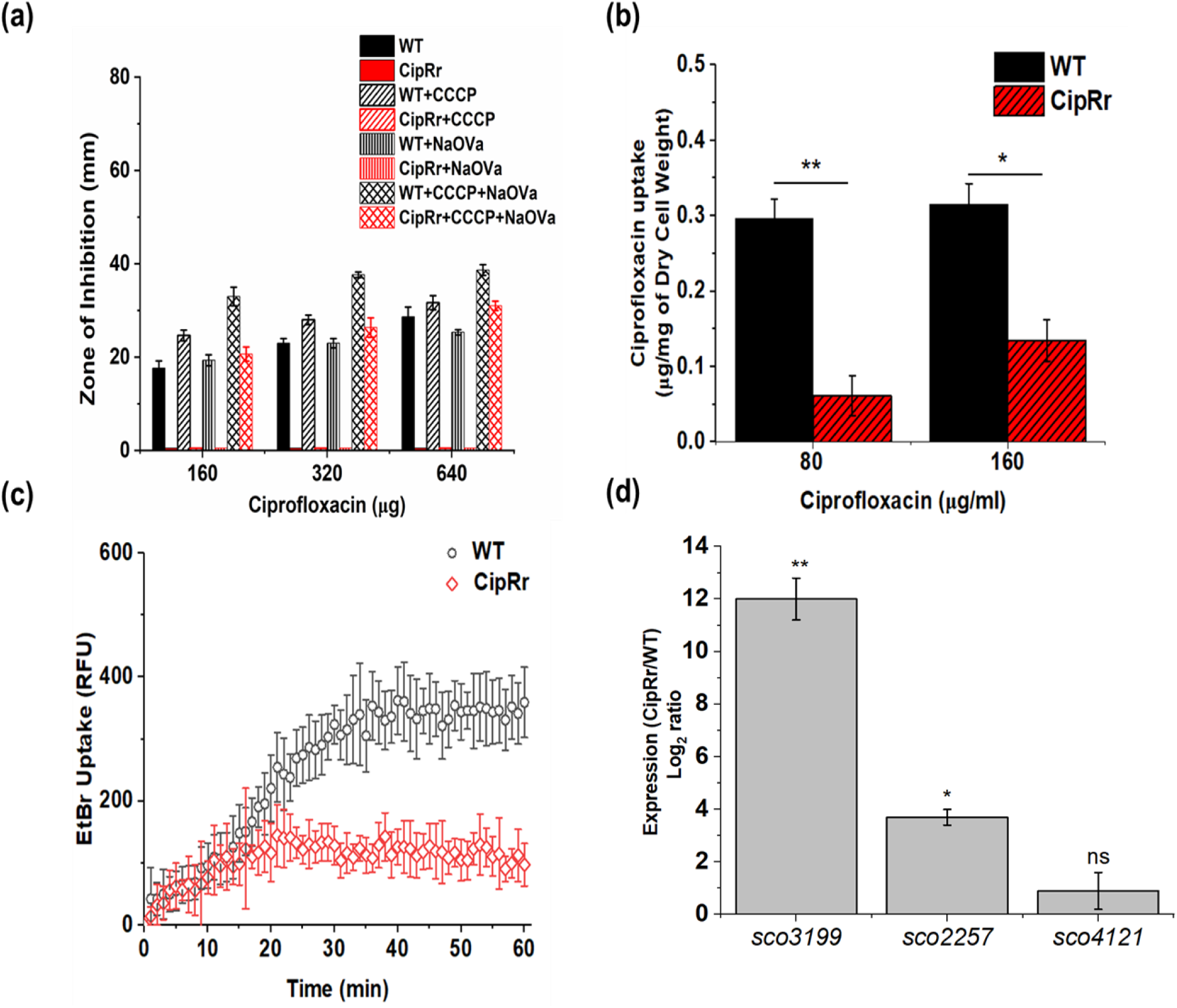
Antibiotic resistance in CipRr is contributed partially via efflux pumps. (a) Determination of susceptibility of WT and CipRr cells towards ciprofloxacin in the presence of CCCP and NaOVa. Susceptibility was measured using the Zone of Inhibition assay. Zone sizes are measured in mm. 5 µg of CCCP and 800 µg of NaOVa were used for the assay. (b) Detection of intracellular ciprofloxacin in WT and CipRr strains. The uptake was carried out at 80 µg/ml and 160 µg/ml of external ciprofloxacin concentration (corresponding to MIC and 2X MIC of WT cells). The treatment was done for 4 hours before measuring the intracellular ciprofloxacin levels. (c) Estimation of EtBr accumulation was carried out in WT and CipRr strains. The accumulation was studied over a period of 60 minutes. EtBr at a concentration of 5µg/ml was used for the assay. (d) Expression of the MFS and the ABC transporters *sco3199* and *sco2257,* respectively measured in CipRr using Real-Time quantitative PCR. The expression was normalized using 23S RNA as the housekeeping gene and further normalized with the WT cells. Expression of the MFS transporter *sco4121* was taken as a negative control. Statistical significance calculated with respect to WT cells.** Indicates p<0.005, * Indicates p<0.05.

To further support this data, we measured the intracellular ciprofloxacin concentration in CipRr and compared it to that of WT cells. We observed a 3-fold lower intracellular ciprofloxacin in CipRr strains as compared to WT when treated with 80 µg/ml and a 2-fold decrease when treated with 160 µg/ml (Figure 4b). We also measured the accumulation of the aromatic compound EtBr, recognized as a substrate by many efflux pumps to monitor overexpression of efflux pumps (24). In our previous studies, we have successfully used EtBr to monitor overexpression of efflux pumps in *S. coelicolor* (13, 25). When recorded over a period of 60 minutes, CipRr displayed nearly 4-fold lower accumulation of EtBr, suggesting the overexpression of efflux pumps in CipRr is related to elevated resistance (Figure 4c). The cell viability remained unaffected under these conditions.

Further, we also observed the expression of *sco3199,* encoding an MFS transporter, and the expression of *sco2257,* encoding an ABC transporter, using qRT-PCR. These transporters were previously found to be upregulated in response to ciprofloxacin stress under different growth conditions (12, 13). As compared to the WT, both *sco3199* and *sco2257* were significantly upregulated to nearly 11-fold (log_2_ ratio) and 3-fold (log_2_ ratio), respectively, in CipRr. This data thus suggests that overexpression of a diverse group of efflux pumps is responsible for the elevated resistance seen in CipRr strains (Figure 4d).

### Whole genome Sequencing identifies distinct mutations associated to CipRr phenotype

Whole Genome Sequencing of CipRr and WT strains revealed a total of 6 unique Single Nucleotide Polymorphisms (SNPs), and 3 unique Insertions and Deletions (InDels). The details of the positions and the mutation type are listed in Table 2. Two non-synonymous SNPs were observed in the coding regions of *sco1614* and *sco3200*, encoding GntR and DeoR family transcriptional regulators, respectively. Both regulator families are known to play essential roles in governing morphological phenotypes, antibiotic production, oxidative stress tolerance, and efflux pump regulation in *Streptomyces* (26–30) *W*e thus hypothesize that these mutations observed in *sco1614* and *sco3200* may be involved in the altered morphology, higher yields of endogenous antibiotic production, increased tolerance to oxidative stress, and increased expression of efflux pumps as seen in CipRr strains. Another SNP was identified in *sco3973*, which encodes a small noncoding RNA. In *S. coelicolor*, several small non-coding RNAs are reportedly important in aerial hyphae development and antibiotic production (31). This mutation can thus additionally contribute to the distinct morphology of the mycelia and increased yield of endogenous antibiotics as noted in CipRr.

**Table 2:**
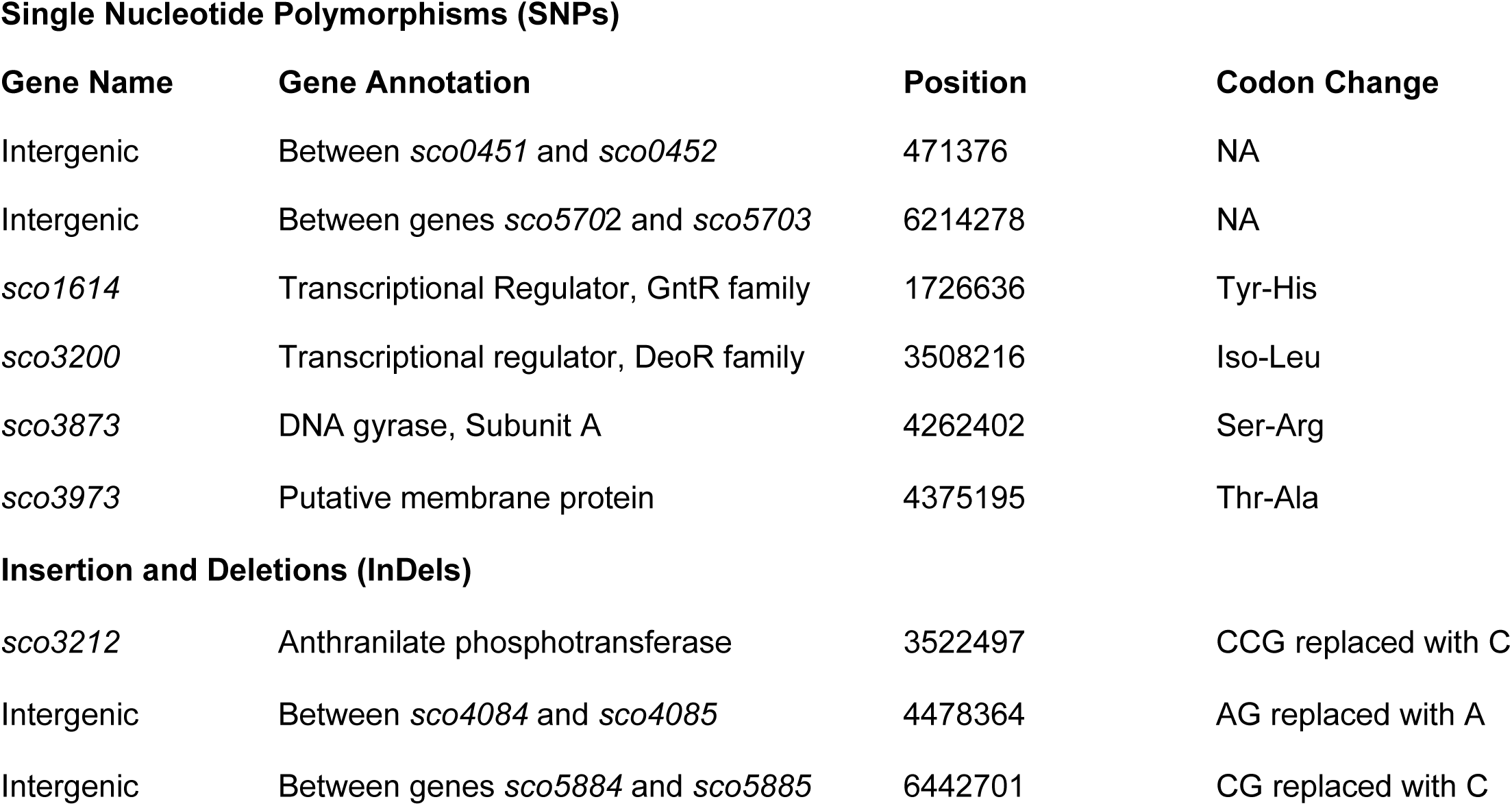
List of Single Nucleotide Polymorphisms (SNPs) and Indels (Insertion and Deletions) observed in CipRr as compared to WT. SNPs and InDels were detected with depth greater than 50 and frequency greater than 50%.

**Single Nucleotide Polymorphisms (SNPs)**
| **Gene Name** | **Gene Annotation** | **Position** | **Codon Change** |
| --- | --- | --- | --- |
| Intergenic | Between *sco0451* and *sco0452* | 471376 | NA |
| Intergenic | Between genes *sco5702* and *sco5703* | 6214278 | NA |
| *sco1614* | Transcriptional Regulator, GntR family | 1726636 | Tyr-His |
| *sco3200* | Transcriptional regulator, DeoR family | 3508216 | Iso-Leu |
| *sco3873* | DNA gyrase, Subunit A | 4262402 | Ser-Arg |
| *sco3973* | Putative membrane protein | 4375195 | Thr-Ala |
| **Insertion and Deletions (InDels)** |  |  |  |
| *sco3212* | Anthranilate phosphotransferase | 3522497 | CCG replaced with C |
| Intergenic | Between *sco4084* and *sco4085* | 4478364 | AG replaced with A |
| Intergenic | Between genes *sco5884* and *sco5885* | 6442701 | CG replaced with C |

CipRr harbored another non-synonymous mutation in *sco3873*, encoding for the A-subunit of DNA gyrase A protein, the target of fluoroquinolones (32). However,, the mutation was located outside the Quinolone Resistance Determining Region (QRDR) (Table 2) and unlikely to confer high-level ciprofloxacin resistance. Additionally, two SNPs were found in the intergenic regions between *sco0451* and *sco0452 (*encoding SIR-like regulatory protein)as well as between *sco05702*(encoding a probable lipoprotein) and *sco5703* (encoding a homolog of Rim-P protein of *E. coli).* Apart from these SNPs, three unique InDels were detected in the CipRr strain, two of these InDels were confined to the intergenic region between *sco4084* and *sco4085*(both encoding hypothetical proteins), and *sco5884* (encodes for a Serine/ Threonine Kinase) and *sco5885*(encodes a membrane protein). The third InDel was present in the coding region of *sco3212*, encoding an anthranilate phosphoribotransferase. These InDels, primarily located in or near metabolic genes, may contribute to the altered metabolite profiles and phenotypes in CipRr strains.

## Discussion

Ciprofloxacin is a broad-spectrum antibiotic commonly used for the treatment of urinary tract infections and infections of the lower respiratory tract, and also as second-line treatment of pulmonary tuberculosis (33, 34). However, its uncontrolled use has resulted in the emergence of resistance in different pathogen rendering the drug ineffective. Studies indicate elevated intrinsic resistance to ciprofloxacin in *Streptomyces* isolated from different geographical areas, implying the presence of uncharacterized resistance determinants (3). Previously, through transcriptomic studies, we have identified several genes that are differentially expressed in response to variable ciprofloxacin stress (12, 13). In the present study, we utilized adaptive laboratory evolution to select for the resistance determinants that may emerge upon prolonged ciprofloxacin exposure. CipRr was generated by serially passaging the WT *S. coelicolor* under increasing ciprofloxacin concentrations, reaching a final 4× MIC (320 µg/ml), resulting in high-level resistance. Beyond ciprofloxacin, CipRr exhibited a cross-resistance to ofloxacin and moxifloxacin but increased sensitivity towards chloramphenicol, tetracycline, and novobiocin. This is consistent with the collateral sensitivity pattern that occurs as a trade-off of the evolutionary pathway, where resistance to one drug makes the cells hypersensitive to others

(15). Another observation was a distinct phenotype characterized by a nodular display of aerial hyphae in CipRr when compared to WT cells. Previous reports have linked antibiotic-induced morphological changes to changes in peptidoglycan synthesis, facilitating survival under stress (35). We hypothesize that CipRr undergoes cell wall rearrangements that enhance tolerance to ciprofloxacin.

In addition to antibiotic resistance, CipRr displayed a significantly elevated production of actinorhodin and undecylprodigiosin. Similar adaptation has been identified in ofloxacin-evolved *S. coelicolor*, where modulations in ACT biosynthetic gene cluster enhanced actinorhodin production (36). Therefore, evolution under increasing ciprofloxacin stress may fix mutations leading to upregulation of biosynthetic genes required for secondary metabolites. Concomitantly, CipRr displayed higher tolerance towards the oxidant HOCl with reduced intracellular ROS generation. This aligns with studies that show induction of oxidative stress response genes due to increased actinorhodin production, aiding cell survival (23). Hence, CipRr strains may adopt tolerance to oxidative stress as a defense mechanism in response to the overproduction of actinorhodin.

Whole genome sequencing (WGS) identified unique mutations associated with differential ciprofloxacin resistance in CipRr. Notably, non-synonymous mutations were determined in the GntR and DeoR group of transcriptional regulators SCO1614 and SCO3200, respectively. GntR and DeoR group of transcriptional regulators are essential in bacterial metabolism and regulation of secondary metabolite synthesis and export in streptomyces (26, 29). For example, the GntR regulator SCO3932 governs actinorhodin biosynthesis by binding to *actII-orf4* promoter, while SCO1728 and SCO7168 are also known to influence actinorhodin production (37). Similarly, the DeoR regulators SdrA and AtrA regulate secondary metabolism in *S. avermitilis* and *S. roseoporus,* respectively (38, 39). Both groups are also known to regulate oxidative stress in other bacteria, like the GntR reglator YtrA in *Xanthomonas citri* (40) and the DeoR regulator FruR in *Listeria monocytogenes* (41). Furthermore, they are also implicated in morphological development, including aerial hyphae formation and differentiation (26, 38).

In addition to these regulators, SNPs and InDels were also noted in the metabolic genes, such as *sco3212*, encoding anthranilate phosphoribotransferase and several intergenic regions possibly contributing to the CipRr phenotype (Table 2). Interestingly, the susceptibility of CipRr to ciprofloxacin could be restored to WT levels when efflux pump inhibitors CCCP and NaOVa were combined, but not when used individually, indicating the involvement of multiple efflux systems. Additionally, the decreased intracellular accumulation of ciprofloxacin and EtBr corroborates the findings. Moreover, *sco3199 and sco2257 were* significantly upregulated in CipRr, suggesting their involvement in high-level resistance. The *SCO3199* gene shares a 46% identity to *actII-2*, the gene encoding actinorhodin exporter in *S. coelicolor*, suggesting a link between the upregulation of *sco3199* to increased yield of actinorhodin seen in CipRr.

The WGS data also identified a mutation in the DNA gyrase A gene, the target for fluoroquinolones, in CipRr, but outside the Quinolone Resistance Determining Region (QRDR)likely exerting a minimal influence on resistance. Our findings suggest that ciprofloxacin adaptation in *S. coelicolor* fixes mutations in transcriptional regulators and metabolic genes, leading to activation of multiple efflux pumps and enhanced drug tolerance. Additionally, the fixed mutations impact the morphological development, secondary metabolite synthesis, and oxidative stress management.

Given that homologs of many of these genes exist in clinical pathogens (data not shown), we predict that horizontal gene transfer from antibiotic producers can drive fluoroquinolone resistance in pathogens. Therefore, artificial evolutionary strategies in antibiotic producers, such as Streptomyces, can help us elucidate the transcriptionally silent genes, dissemination of which may result in elevated drug resistance in different clinical pathogens. Future transcriptomics analyses of CipRr will further elucidate the role of the fixed mutations in modulating antibiotic resistance, morphological development, and secondary metabolite production.

## Materials and Methods

### Bacterial strains, growth media, and conditions

Spores of *Streptomyces coelicolor* (WT and CipRr) were germinated in 2XYT broth (Tryptone-1.6%, NaCl-1%, Yeast Extract-1%) for 8 hours. The germinated mycelia were further inoculated into R5 media (10%Sucrose, 1% Glucose, 0.5% Yeast Extract, 1% MgCl_2_.7H2O, 0.5% TES buffer, 0.01% Casein Hydrolysate, 0.01% Potassium phosphate (K_2_SO_4_)) and grown at 30°C for all assays. For growing spores of *S. coelicolor*, Mannitol Soya Agar (Mannitol-%, Soya-2%, Agar-2%) plates were used. List of strains mentioned in Table 3.

**Table 3:** List of strains used in this study.

| Strains | Features | Reference |
| --- | --- | --- |
| *S. coelicolor* |  |  |
| WT M145 | *S. coelicolor* WT cells evolved in the absence of any drug | (25) |
| CipRr | *S. coelicolor* ciprofloxacin resistant strain evolved under increasing ciprofloxacin concentration. | This study |

### Creation of CipRr mutants

Ciprofloxacin-resistant mutants CipRr were created by serially passaging *S. coelicolor* WT strain on R5 agar with gradually increasing concentrations, starting at the MIC (80µg/ml) and reaching 4X MIC (320 µg /ml). Intermittent drug-free passages on R5 plates was carried out to select for stable mutations. All assays were done using the endpoint culture.

### Gene Expression Studies

RNA was isolated from *S. coelicolor* using the Trizol method described previously (42). This was followed by cDNA synthesis using Revert Aid H minus Reverse Transcriptase (RT) enzyme (Thermo Scientific) following the manufacturer’s recommendations. The gene Expression studies using qPCR were carried out on a BioRad CFX96 instrument using SYBR Green iTaq Mix, 0.5 µM each of specific primers (Table 4) and 100ng template. Relative gene expression was calculated by 2^-ΔΔCt^ method, normalized against the housekeeping gene and the untreated control.

**Table 4:**
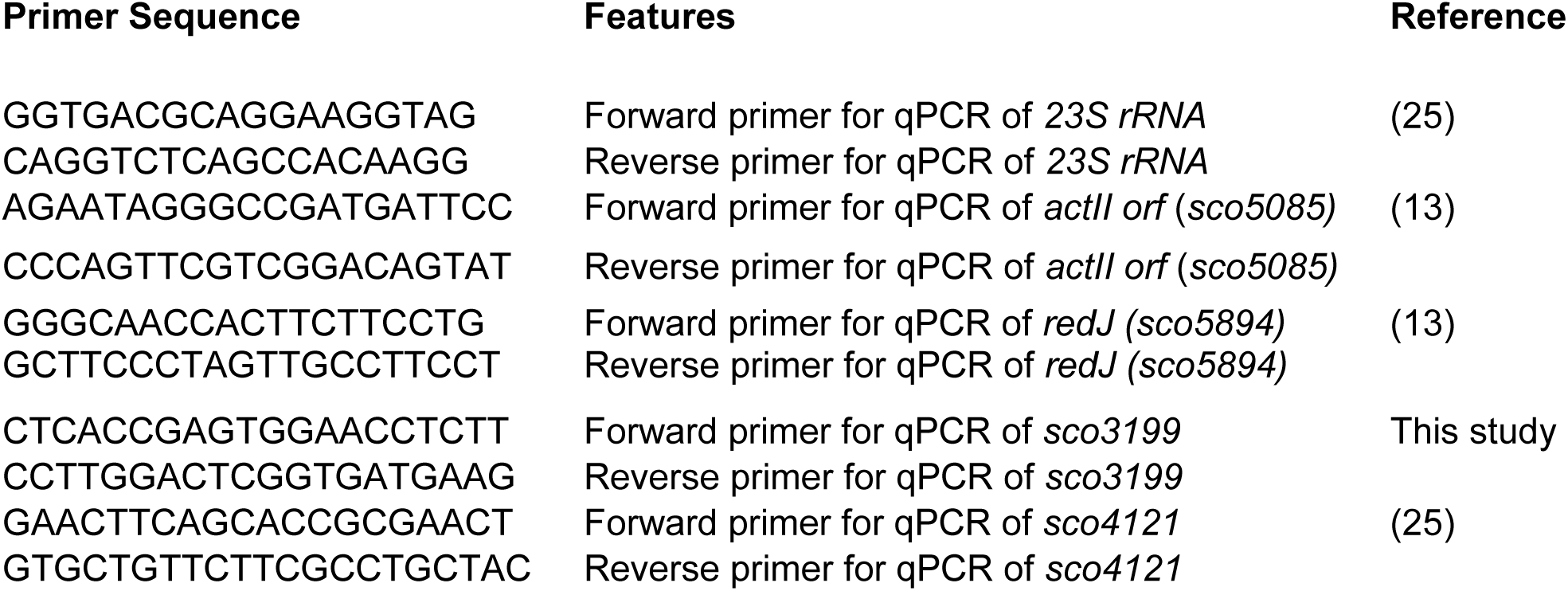
List of Primers used in this study.

### Antimicrobial susceptibility assay

#### (i) Disc Diffusion method

A modified protocol was adopted for measuring drug sensitivity via the Disc Diffusion technique, as documented previously (43). Briefly, 10^8^ spores of the WT and CipRr strains of S. *coelicolor* strains were spread onto R5 plates. Sterile discs impregnated with the desired amount of the drug were placed onto the plates, incubated for 3 days at 30°C and the zone of inhibition around the disc was measured (in millimeters). Increased zone diameter suggests increased sensitivity towards the particular drug. Drugs that were tested include ciprofloxacin (Sigma Aldrich), ofloxacin, moxifloxacin, chloramphenicol, tetracycline, novobiocin, spectinomycin, oleandomycin, EtBr and CCCP (All from Himedia). Oxidant sensitivity was similarly assessed using H_2_O_2_ and HOCl.

#### (ii) Agar Dilution method

Agar dilution method was also used to determine the sensitivity of the strains against various antibiotics, performed with slight modifications (44). R5 agar plates containing the desired antibiotic concentration were prepared and 10^8^ spores of *S. coelicolor* (WT and CipRr) were streaked onto these plates. The plates were incubated for 3 days at 30°C and the growth was monitored and recorded to determine the MIC. The antibiotics tested and the drug sources were consistent across both assays.

### Estimation of membrane permeability by NPN assay

Membrane permeability was assessed using1-Phenylnapthalmine (NPN) dye (18). *S. coelicolor* cells (WT and CipRr) were grown to OD_600_ = 1 at 30°C at 150 rpm. Cells were then harvested and resuspended in 5 mM 4-(2-hydroxyethyl)-1 piperazineethanesulfonic acid (HEPES) buffer. Each sample (100 µl) was mixed with 100 µl of 50 µM NPN and the fluorescence was recorded at 355 nm excitation and 402 nm emission wavelengths.

### Ciprofloxacin Uptake

Accumulation of ciprofloxacin in *S. coelicolor* and CipRr was estimated using a previously defined protocol (25). WT and CipRr cells were grown to OD_600_ = 1, concentrated to OD_600_ = 10and equilibrated at 37 °C for 10 minutes. Ciprofloxacin was added at 80 µg/ml (MIC) and 160 µg/ml (2X MIC) and incubated at 37°C for 4 hours. Following which the cells were chilled, washed with chilled 0.5 M PBS and lysed overnight in 0.1 M glycine (acidified to pH 3 with HCl) kept at 25°C. The intracellular drug uptake was measured by fluorescence at 280 nm excitation and 420 nm emission wavelengths (Molecular Devices). The final uptake of ciprofloxacin was normalized to dry cell weight.

### ROS measurement via DCFDA assay

Intracellular ROS was quantified using H2-DCFDA (2, 7-dichlorofluorescein diacetate), a cell permeable fluorogenic dye. H2-DCFDA upon entering the cells via diffusion, gets deacetylated by the cellular esterases into a non-fluorescent moiety which later is oxidized by the ROS into a fluorescent compound 2, 7, dichlorofluorescein that can be measured spectrophotometrically (45). WT and CipRr were grown to OD_600_ = 1, incubated with 10 µM of DCFDA for 10 minutes, washed and resuspended in 0.1 M PBS. The generation of ROS was monitored for 1 hour at 435 nm excitation and 530 nm emission wavelengths.

### Quantification of Undecylprodigiosin (RED)

Undecylprodigiosin production was quantified as previously described (46). Briefly, WT and CipRr cultures were grown in R5 medium at 30°C. Samples were collected at regular time intervals of 24, 36, 48, 72 and 96 hours, vacuum dried, and extracted with methanol (acidified with HCl to 0.5 M). Absorbance of the supernatant was measured at 530 nm (UV-Vis spectrophotometer), and concentrations were calculated using the Beer-Lambert equation (ε₅₃₀ = 100,500).

### Quantification of Actinorhodin (ACT)

Actinorhodin was quantified as per the published protocol (47). Briefly, WT and CipRr cultures were grown in R5 media at 30°C. Samples were collected at regular time intervals of 24, 36, 48, 72 and 96 hours, lysed with equal volume of 0.1N KOH and the cellular debris removed by centrifugation at 12,000 rpm for 10 minutes. Absorbance at 640 nm was used to calculate actinorhodin concentrations (ε₆₄₀ = 25,320) using Beer Lambert equation.

### Cryo SEM Imaging

Cells for microscopy were prepared as described previously (12). The cells were harvested through centrifugation and re-suspended in a 50 mM sodium phosphate buffer (pH 7.0) to achieve OD_600_ = 10 and incubated at 30 °C for 10 minutes For the treatment phase, ciprofloxacin was introduced into the culture flasks at final concentrations of either 20 μg/ml or 80 μg/ml, while a control flask was maintained without any treatment followed by incubation for 4 hours. After the incubation period, 100 µl of samples were taken from each flask, adjusted to approximately 10^6 cells/ml and imaged using phase contrast microscope (Nikon Eclipse E 200 at × 40) and the CRYO-FEG-SEM (JSM-7600F) platform.

### DNA extraction and Whole Genome Sequencing

*S. coelicolor* WT and CipRr were grown in liquid R5 media to OD_600_ = 1. Cells were pelleted by centrifugation at 6000rpm for 10 minutes, resuspended in deionized water and digested with lysozyme (1 mg/ml) followed by incubation at 37°C for 4 hours. Genomic DNA was isolated using standard phenol-chloroform extraction, followed by isopropanol wash and (100%) ethanol precipitation at -80°C. DNA pellets were washed with 70% ethanol, resuspended in double-distilled water and treated with RNAse (50 µg/ml). The isolated DNA was quantified and paired-end sequencing (150 bp) was performed by Beijing Genomics Institute (BGI, Hong Kong) using Illumina Hiseq Xten.

### Sequence Analysis

The sequence reads were aligned to the *S. coelicolor* reference sequence retrieved from NCBI (NC_000388.3) using the Bowtie 2 alignment in Galaxy. PCR duplicates were removed (Mark Duplicates) and read statistics were generated using FLAGSTAT. The variants were identified with VarScan with ≥50× coverage and ≥50% variant frequency. Unique mutations in CipRr were compared to the WT genome, which was sequenced along with the CipRr strain.

## Acknowledgement

We acknowledge the Department of Chemical Engineering, IIT Bombay, for the CRYO-FEG-SEM facility.

## Data Availability

The data has been submitted to NCBI Sequencing Read Archive with the Bioproject ID PRJNA1391826.

